# A novel mathematical method for disclosing oscillations in gene transcription: a comparative study

**DOI:** 10.1101/151720

**Authors:** Athanasios C. Antoulas, Bokai Zhu, Qiang Zhang, Brian York, Bert W. O’Malley, Clifford C. Dacso

## Abstract

Circadian rhythmicity, the 24-hour cycle responsive to light and dark, is determined by periodic oscillations in gene transcription. This phenomenon has broad ramifications in physiologic function. Recent work has disclosed more cycles in gene transcription, and to the uncovering of these we apply a novel signal processing methodology known as the pencil method and compare it to conventional parametric, nonparametric, and statistical methods. Methods: In order to assess periodicity of gene expression over time, we analyzed a database derived from livers of mice entrained to a 12-hour light/12-hour dark cycle. We also analyzed artificially generated signals to identify differences between the pencil decomposition and other alternative methods.

**Results:** The pencil decomposition revealed hitherto-unsuspected oscillations in gene transcription with 12-hour periodicity. The pencil method was robust in detecting the 24-hour circadian cycle that was known to exist, as well as confirming the existence of shorter-period oscillations. A key consequence of this approach is that orthogonality of the different oscillatory components can be demonstrated. thus indicating a biological independence of these oscillations, that has been subsequently confirmed empirically by knocking out the gene responsible for the 24-hour clock.

**Conclusion:** System identification techniques can be applied to biological systems and can uncover important characteristics that may elude visual inspection of the data. Significance: The pencil method provides new insights on the essence of gene expression and discloses a wide variety of oscillations in addition to the well-studied circadian pattern. This insight opens the door to the study of novel mechanisms by which oscillatory gene expression signals exert their regulatory effect on cells to influence human diseases.

## Introduction

Gene transcription is the process by which the genetic code residing in DNA is transferred to RNA in the nucleus as the inauguration of protein synthesis. The latter process is called translation and occurs in the cytoplasm of the cell. Circadian rhythm, the 24-hour cycle that governs many functions of the cell, is the result of a complex interaction of transcriptional and translational processes. The importance of circadian rhythm to physiologic processes has been underscored in 2017 by the awarding of the Nobel Prize in Physiology or Medicine to the investigators who described the molecular mechanisms controlling it. However, in addition to the circadian oscillation driven by light and dark, other so-called infradian and ultradian rhythms have clear biologic import. Blood pressure, some circulating hormones, and some physiological functions appear to have 12-hour periodicity whereas other processes such as the menstrual cycle more closely follow a lunar cycle.

Accordingly, we sought to uncover novel 12-hour oscillations in gene expression. In many cases, the 12-hour gene oscillation is superimposed on the 24-hour cycle; thus it is hidden in conventional analysis. Additionally, experiments designed to elucidate the 24-hour circadian often do not have the granularity required to reveal an interval of less than 24 hours as they are constrained by the Shannon-Nyquist Sampling Theorem [1].

To reveal periodicities in gene expression other than the 24-hour circadian cycle, we applied digital signal processing methodology to this biologic phenomenon. Although this approach is, to our knowledge, less commonly used in the biological field, it is justified because the transcription of DNA to RNA is indeed a signal, packed with information for making the enormous repertoire of proteins.

To extract the fundamental oscillations (amplitude and period) present in the data, we utilized publicly available time-series microarray datasets on circadian gene expression in mouse liver (under constant darkness) [2] and analyzed over 18,000 genes spanning a variety of cellular process ranging from core clock control, metabolism, and cell cycle to the unfolded protein responses (UPR), a measure of cell stress. In addition, one set of measurements of RER (respiratory exchange ratio) from wild-type mice (generated by us) was also performed. We constructed linear, discrete-time, time-invariant models of low order, driven by initial conditions, which approximately fit the data and thus reveal the fundamental oscillations present in each data set. In addition to the 24-hour (circadian) cycle known to be present, other fundamental oscillations have been revealed using our approach.

## Methods

We searched for 12-hour oscillations in several biological systems. Systems were chosen that represented not only gene transcription but also phenotype; they represent the way in which these biological systems are expressed in the whole organism. The reasoning was that if the 12-hour oscillation in transcription was biologically significant, it would be represented in some measurable function of the cell.

Initially, we analyzed a set of transcription data [2] that was collected in mouse liver obtained from animals in constant darkness after being entrained in a 12-hour light/12-hour dark environment. Mice were sacrificed at 1-hour intervals for 48 hours, thus providing enough data points to analyze the signal. The dataset thus obtained contains RNA values for all coding genes. The RNA data were generated using a standard microarray methodology. In addition, RER (respiratory exchange ratio) measurements in mice were also measured and analyzed. The novelty in our analysis consists in using the so-called matrix-pencil method [3]. This is a data-driven system-identification method. It constructs dynamical systems based on time-series data and finds the dominant oscillations present in the ultradian or infradian rhythms. Our purpose here is to compare this method with other established strategies for spectral estimation, including both parametric spectrum estimation methods like MUSIC (MUltiple Signal Classification), ESPRIT (Estimation of Signal Parameters via Rotational Invariance Techniques), and Prony’s (least squares) as well as classical nonparametric models like wavelet transforms and statistical methods like RAIN. These are compared with each other using both artificial and measured data.

### Basic signal processing methods

- **The data**. We consider finite records of data resulting as described above. Generically they are denoted by **y** _*i*_, *i* = 1, *…, N*.
- **Basic model: sum of exponentials**. We seek to approximate the data by means of linear combinations of exponentials plus noise. Thus we seek *k* pairs of complex numbers *α*_*i*_, *β*_*i*_, *i* = 1, 2, *…, k*, such that

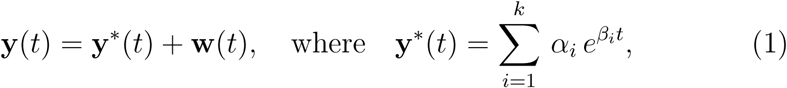

is the noiseless part of the signal and **w**(*t*) is the noise. The requirement is: **y**(*m*) ≈ **y**_*m*_, *m* = 1, 2, *…, N*. Existing approaches to address this problem are MUSIC, ESPRIT, Prony’s (least squares) method, wavelet transform and statistical methods described later.
- **Second model: descriptor representation**. The equivalent *descriptor* model uses an associated *internal variable* **x**(*t*) *∈* ℝ^*k*^ of the system. The resulting equations are:

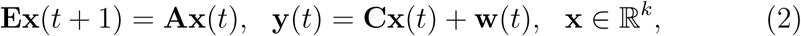

with initial condition **x**(0) = **x**_0_ *∈* ℝ^*k*^, where **E**, **A** *∈* ℝ^*k×k*^, **C** *∈* R^1*×k*^.
- **Third model: AR (Auto Regressive) representation**. The above model can also be expressed as an AR model driven by an initial condition. As above we let **y**(*t*) = **y\*** (*t*) + **w**(*t*), (where **y\*** (*t*) is the noiseless term and **w**(*t*) the noise). It follows that (1) can be rewritten as:

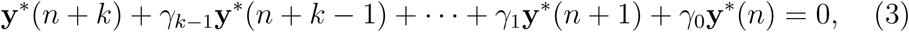

with initial conditions **y\*** (𝓁), 𝓁= 0, 1, *…, k -* 1.

**Goal**. Discover the fundamental oscillations inherent in the gene data, using these models and reduced versions thereof.

### Processing of the data with the pencil method

The data **y**_1_, **y**_2_, …, **y**_*N*_, are used to form the *Hankel matrix*:

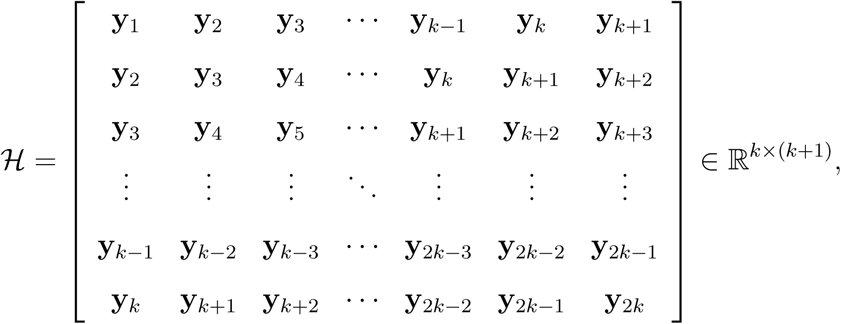

where for simplicity it is assumed that *N* = 2*k*. Then we define the quadruple (**E**, **A**, **B**, **C**):

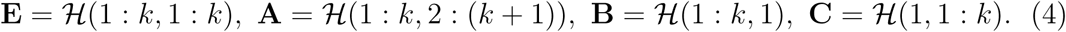

This quadruple constitutes the *raw model* of the data. This model is linear, time-invariant and discrete-time with a non-zero initial condition:

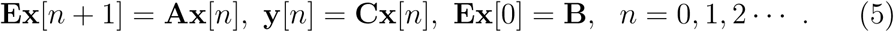

#### Reduced models and fundamental oscillations

The **dominant** part of the raw system is determined using a **model reduction** approach [4], [5], [6], [3]. The procedure is as follows.

**Pencil procedure for obtaining dominant sub-models**

- Compute the SVDs:

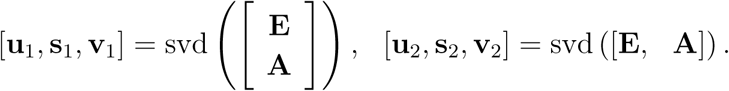
- Choose the dimension *r* of the reduced system (e.g *r* = 3, *r* = 5, *r* = 7 etc.). Then

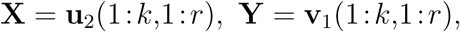

are used to project the raw system to the subdominant system of order *r*:

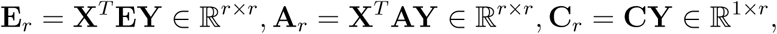

and **x**_*r*_ = **X**^*T*^ **x**_0_ *∈* R^*r*×1^.

The associated reduced model of size *r* is then:

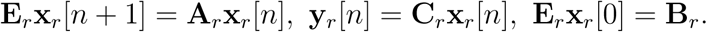

Assuming (as is usually the case) that **E**_*r*_ is invertible, the approximated data can be expressed as:

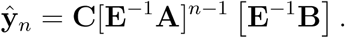

#### Estimating *r*

Important byproducts of the pencil method are the singular values **s**_1_ and **s**_2_ mentioned above. The accuracy of the approximation is determined by the first neglected singular singular value *σ*_*r*+1_, as the resulting approximation error is proportional to this singular value. This implies the following rule.

#### Rule

choose *r* so that 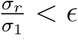, where *∊* is a tolerance which depends on the data at hand. For instance ϵ = 0.01, implies roughly speaking that data contributing less than 1% to the overall result are discarded. In this regard the following remark is in order. The data considered in this paper are rather short-duration and therefore in many cases we have not truncated the data.

#### Partial fraction expansion of the associated transfer function

**H**_*r*_(*z*) = **C**_*r*_(*z***E**_*r*_*-* **A**_*r*_)^-1^**B**_*r*_. This involves the eigenvalue decomposition (EVD) of the matrix pencil (**A**_*r*_, **E**_*r*_), or equivalently of 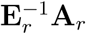; let

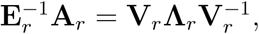

where the columns of **V**_*r*_ = [**v**_1_, …, **v**_*r*_] are the eigenvectors, **Λ**_*r*_ = diag[*λ*_1_, *…, λ*_*r*_] are the eigenvalues of the reduced system (poles of **H**_*r*_(*z*)), and 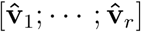 are the rows of 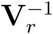. The approximate data can be expressed as:

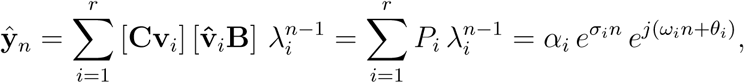

where 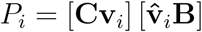, is the complex amplitude of the *i*^*th*^, oscillation; expressing this in polar form 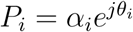, *α*_*i*_ is the real amplitude and *θ*_*i*_ the phase. Finally, if we express the eigenvalues as 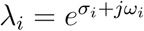, *σ*_*i*_ is the decay (growth) rate, and *ω*_*i*_ the frequency, of the *i*^th^ oscillation.

#### Poles and oscillations

Often in (digital) system theory, the quantity λ_*i*_*∈* ℂ is referred to as *pole* of the associated system. Oscillatory signals result when *σ*_*i*_ = 0, which it turn implies that the magnitude of the pole λ_*i*_ is equal to one: *│ λ*_*i*_*│*= 1, and the period of oscillation is 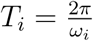. For instance a signal with λ_*i*_ = 1, represents a constant (step), while signals with 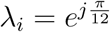,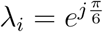 (which are both on the unit circle with angles 15^*°*^, 30^*°*^ degrees) represent pure oscillatory signals with periods 24, 12 hours respectively.

#### Angle between signals and orthogonality

In the sequel we will make use of angles between signals. Here we briefly define these concepts. Given discrete-time finite duration signals (vectors)

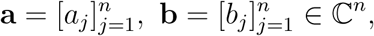

their **inner product** is defined as

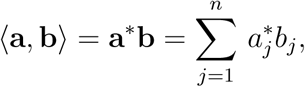

where (**·**)* denotes complex conjugation and transposition; the **angle** between these signals is defined as

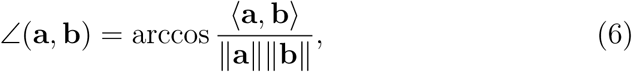

where ‖ **·** ‖ denotes the Euclidean 2-norm. **Orthogonality** means that the angle between the two signals is 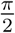, or equivalently that their inner product is zero; this is sometimes denoted by **a** ⊥ **b**. In the sequel we also make use of the symbol 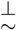 to indicate *approximate orthogonality*, i.e. an angle between signals close to 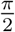 radians or 90^*°*^ degrees.

### Other methods

To complete the picture, we briefly list other methods which can be used to analyze the gene data.

## MUSIC

The MUSIC algorithm [7], [8], is a parametric spectral estimation method based on eigenvalue analysis of a correlation matrix. It uses the orthogonality of the signal subspace and the noise subspace to estimate the frequency of each oscillation. It assumes that a set of data can be modeled as **Y** = **Γa** + **n**, where **Y** = [**y**_1_ **y**_2_ *…* **y**_*N*_]^*T*^*∈*ℜ^*N*^, is a set of gene transcription data, **Γ** = [**e**(*ω*_1_) **e**(*ω*_2_) **e**(*ω*_*K*_)] is the transpose of a Vandermonde matrix, *K* is the number of dominant frequencies, and **e**(*ω*_*i*_) = [1 *e*^*jω*^_*i*_*… e*^*j*(*K*–1)*ω*^_*i*_]^*T*^, **a** = [*a*_1_ *a*_2_ *… a*_*K*_]^*T*^ contains the amplitudes of the dominant *K* frequencies, 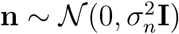, is white noise. The autocorrelation matrix is

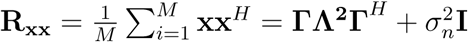

where **Λ** = diag(λ_*i*_) and *M* is the number of columns in the Hankel matrix. We can see that the rank of matrix **ΓΛ^2^Γ**^*H*^ equals *K* where the nonzero eigenvalues are 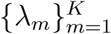. Then the sorted eigenvalues of the autocorrelation matrix **R_xx_** can be expressed as

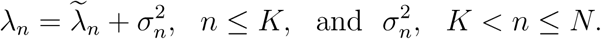

It follows that the noise subspace contains the eigenvectors of the autocorrelation matrix **R_xx_** corresponding to the *N − K* smallest eigenvalues. Then

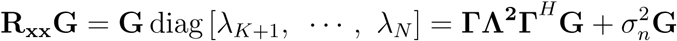

so **Γ**^*H*^**G** = 0, and the frequency values 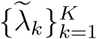 are the only solutions of **e**(*ω*)^*H*^**GG**^*H*^**e**(*ω*) = 0. The MUSIC algorithm seeks the peaks of the function 1*/*[**e**(*ω*)^*H*^**GG**^*H*^**e**(*ω*)], where *ω ∈* [0, 2*π*]. The Root MUSIC algorithm seeks the roots of **p**^*H*^(*z*^-1^)**GG**^*H*^**p**(*z*) that is the Z-transform of **e**(*ω*)^*H*^**GG**^*H*^**e**(*ω*) where *z* = *e*^*jω*^ ∈ C.

The MUSIC algorithm can only provide the frequency information of the signal. To obtain the amplitude of each oscillation, we need to apply least squares fitting, where the amplitudes of dominant oscillations satisfy **a** = (**Γ**^*H*^**Γ**)^-1^**Γ**^*H*^**x**. It should mentioned that in contrast with the pencil method, MUSIC cannot provide the decay (growth) rate of the oscillations.

## ESPRIT

This is another parametric spectral estimation algorithm [7], [8]. It analyzes the subspaces of the correlation matrix. It estimates the poles relying on rotational transformation. As in MUSIC: 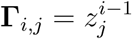, *j* = 1, *…, K, i* = 1, *…, N*, where *z*_*j*_ are the poles. We can construct **Γ**_1_ = **Γ**(1: *N -* 1,:), and **Γ**_2_ = **Γ**(2: *N,*:). The relationship between these two quantities is **Γ**_2_ = **Γ**_1_**Ф**, where **Ф** = diag [*z*_1_, *z*_2_, …, *z*_*K*_],, is the phase shift matrix that represents a rotation. Now we construct a similar structure applying on signal subspace **S** that contains the eigenvectors of the autocorrelation matrix **R_xx_** corresponding to the *K* largest eigenvalues. Let

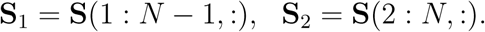

Note that the relationship between **S**_1_ and **S**_2_ is **S**_2_ = **S**_1_**Ψ**. Because **Γ** and **S** have the same column space (see [7, 8]), we have that **Γ** = **ST**, where **T** is an invertible subspace rotation matrix. So we have **Ψ** = **T**^-1^**ФT**. Therefore the poles are the eigenvalues of **Ψ**. Finally least square (LS) to obtain 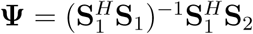. The eigenvalues of **Ψ**, are the poles *z*_*i*_ = *e*^*jωi+σi*^. Thus ESPRIT can estimate both the frequency and the decay (growth) rate of the oscillations. However, as with MUSIC, we need to use LS to obtain the amplitude of each oscillation.

### Wavelet transform

Wavelet transforms can be divided into two categories, the continuous (CWT) and the discrete (DWT) versions. CWT is more suitable for analyzing biologic rhythms because of the associated heat maps are two-dimensional.

In CWT a time signal *x*(*t*) is convolved with a wavelet function. This leads to a time-frequency representation which provides spectrum information in a local time window. This transform can be expressed as 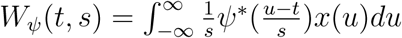, where *s* is the frequency scale, *ψ\** (*t*) is the wavelet function. Since the signal data is obtained by sampling, we can approximately rewrite the equation as 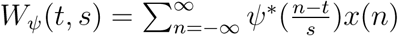. It follows that the integral or sum is applied on the range *-∞* to *∞* that means the domain of signal *x*(*t*) or *x*(*n*) should be the range from *-∞* to *∞*. But the signals considered have finite length, in which case the edge effects become obvious, especially in the low-frequencies.

In practice, there are many wavelet functions that can be chosen, both real- and complex-valued. Real-valued wavelets are useful for treating peaks and discontinuities of signals while complex-valued wavelets yield the information of amplitude and phase simultaneously [9].

### Statistical methods

In this section three statistical methods, namely ARSER, JTK CYCLE and RAIN, will be investigated and their ability to detect biological rhythms evaluated. Those methods focus on the (one) most dominant oscillation in the data, especial JTK CYCLE and RAIN. These constitute statistical tests that calculate the p-value to determine whether a certain rhythm exists in the data [10–12].

## ARSER

ARSER uses the autoregressive (AR) model to obtain the period of oscillation. It then uses linear regression (harmonic) to determine the amplitude and the phase of the oscillation. Finally applying the F-test to pre-processed data and regressive data determines whether an oscillation exists.

### Pre-processing the data

Because the data may not be stable, ARSER applies linear detrending to the raw data. It then uses linear regression to fit the data as a straight line. Subsequently ARSER uses a fourth-order Savizky-Golay algorithm to smooth the data. This low-pass filter removes the pseudo-peaks in the spectrum.

### Finding the period

ARSER uses an autoregressive model to get the period of the oscillation. Given a pre-processed dataset 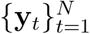 with period interval Δ.

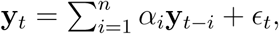

where ϵ_*t*_ is white noise, *α*_*i*_ are AR coefficients, *n* is the order of model (we choose *n* =length-of-data/Δ). To calculate the coefficients, ARSER uses the Yule-Walker method, maximum likelihood estimation and the Burg algorithm. After AR modeling, ARSER can calculate the spectrum:

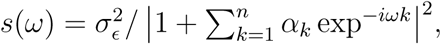

where 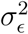 is the variance of white noise. ARSER finds the peaks in time window *t* ∈ [20, 28] as the periods {*T*_*i*_} the oscillation (the optimal periods are determined by Akaike’s information criterion).

### Harmonic Regression

Now we can express the pre-processed data as:

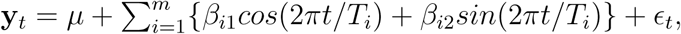

where *β*_*i*1_ and *β*_*i*2_ are the amplitudes. ARSER calculates those amplitude through linear regression.

### F-test

Using the F-test compares the approximation data 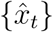 and pre-processed data {*x*_*t*_}. The null and the alternative hypotheses are respectively

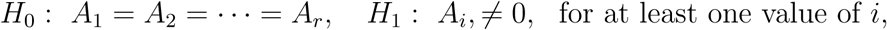

where *A*_*i*_ are the amplitudes which are calculated using linear regression, and *r* is the number of coefficients obtained by linear regression. We can calculate the F coefficient by:

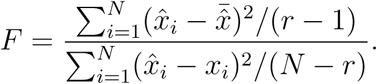

Then we can calculate the p value using the F-distribution *p* = *P* (*F, r* - 1, *N - r*), where *P* (·) is the probability function used to calculate the *p* value based on *F*-distribution.

## JTK_CYCLE and RAIN

JTK CYCLE and RAIN use statistical method to detect the trend in data. The former can find the increasing or decreasing trend in data and RAIN is a development of JTK CYCLE which can combine these two.

A periodic waveform should start from the trough and increase to the peak following a decreasing part to a new trough. Because our data is sampling from the waveform, we can regard every time sampling data point as a variable. Thus we can get *n* variables 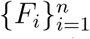 for the waveform such that *T* = *n*Δ (*T* is the period of the waveform, Δ is the time interval of sampling point). We assume the variances of those variables are the same. And they have the same mean value only when the data only have noise without periodic oscillation. So the null and the alternative hypotheses are *H*_0_: *F*_1_ = *F*_2_ = *…* = *F*_*n*_,*H*_1_: *F*_1_ *< F*_2_ *< … < F*_*n*_ or *F*_1_ *> F*_2_ *> … > F*_*n*_. The alternative hypotheses for RAIN is

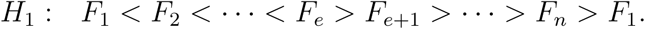

### Calculating the statistical coefficient of trend

Every variable *F*_*i*_, corresponds to a sampling dataset 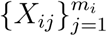, where *m*_*i*_ is the number of sampling data point of the *i*^*th*^ variable 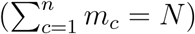. Let 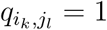, if *X*_*ik*_*≤ X*_*jl*_, and 0 otherwise, and 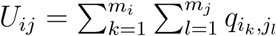 which is the Mann-Whitney U-statistic for comparison of two variables. For JTK CYCLE, the statistical coefficient of trend is

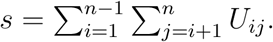

For RAIN, the statistical coefficient of trend is

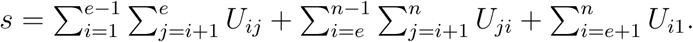

### Calculating the p-value

For the test, the p-value 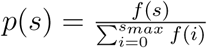. In order to calculate the p-value, we should make clear the distribution *f* (*i*) of statistical coefficient *s* when the null hypotheses *H*_0_ is true. Furthermore the distribution *f* (*i*) is computed, using a generating function 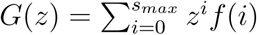. For JTK_CYCLE and RAIN we have respectively:

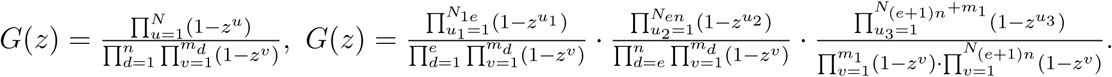

Thus *G*(*z*) for JTK̲CYCLE and RAIN are both polynomials. We can get the distribution *f* (*i*) by calculating the coefficients of *G*(*z*), which can be used in the p-value equation.

## Experimental Results: artificial data

In this section we test the performance of different methods using artificially generated signals. For the continuous wavelet transform, we chose the *complex morlet wavelet* because it allows changes to the resolution in frequency and time domain. For simulation data, we assume the data has the form

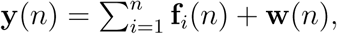

where **w** is white noise with zero mean and variance *σ*^2^ and **f**_*i*_ is the *i*^*th*^ oscillation, where:

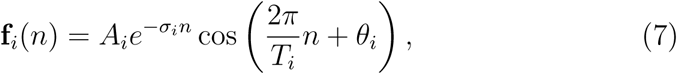

where *A*_*i*_ is the amplitude, *σ*_*i*_ is the decay (growth) rate, *θ*_*i*_ is the phase and *T*_*i*_ is the period. At first we assume that the samples are collected in unit time intervals. The parameters are defined in the table below; the first oscillation is almost constant with small decay; the other three oscillations have a period of approximately 24-12- and 8-hours (see Table 1).

**Table 1.** Parameters used for the simulation

| i | A | $\sigma$ | $\theta$ | T |
| --- | --- | --- | --- | --- |
| 1 | 1 | 0.005 | 0 | ∞ |
| 2 | 1 | 0.004 | $\frac{\pi}{2} - 6$ | 24.8 |
| 3 | 0.3 | -0.002 | $\frac{\pi}{2}$ | 11.8 |
| 4 | 0.1 | 0.005 | $\frac{\pi}{2} + 1$ | 7.5 |

The experiment has the following parts. First, the sensitivity to noise is investigated. Here, the variance of noise is changed and the performance of each of the different methods is examined. Second, the impact of the length of the data is investigated. Finally, the frequency of data collection (can be referred to as *sampling frequency*) will be examined. Recall that the Nyquist sampling theorem provides the lower bound for the sampling frequency in order to prevent aliasing. This can be used to determine appropriate sampling frequencies for continuous-time signals.

### Sensitivity to noise

To test the sensitivity of these various methods to noise, we set the standard deviation of **w** as *σ* = [0, 0.03, 0.1, 0.3].

**Figure 1**

**Curves for simulation data**

This figure shows curves of different methods and simulation data (length 50) with *σ* as stated. The red points are simulation data, blue, green and magenta are the curves of the pencil, ESPRIT and MUSIC methods respectively. This figure shows that the pencil and ESPRIT methods yield a perfect fit in all situations. The MUSIC algorithm gives a good fit only for small amounts of noise. In Table 2, we display the poles obtained by using each method.

**Table 2.** Poles determined by different methods

| $\sigma = 0.01$ | | | | $\sigma = 0.1$ | | | |
| --- | --- | --- | --- | --- | --- | --- | --- |
| orig. poles | Pencil | ESPRIT | MUSIC | orig. poles | Pencil | ESPRIT | MUSIC |
| 0.990 | 0.990 | 0.990 | 1.000 | 0.990 | 0.989 | 0.989 | 1.000 |
| $0.958 \pm 0.248i$ | $0.958 \pm 0.248i$ | $0.958 \pm 0.248i$ | $0.970 \pm 0.239i$ | $0.958 \pm 0.248i$ | $0.960 \pm 0.248i$ | $0.960 \pm 0.249i$ | $0.974 \pm 0.225i$ |
| $0.870 \pm 0.502i$ | $0.870 \pm 0.512i$ | $0.870 \pm 0.512i$ | $0.867 \pm 0.497i$ | $0.870 \pm 0.502i$ | $0.867 \pm 0.511i$ | $0.867 \pm 0.512i$ | $0.834 \pm 0.551i$ |
| $0.662 \pm 0.735i$ | $0.662 \pm 0.735i$ | $0.662 \pm 0.735i$ | $0.693 \pm 0.721i$ | $0.662 \pm 0.735i$ | $0.669 - 0.772i$ | $0.662 \pm 0.751i$ | $-0.974 \pm 0.2235i$ |
| $\sigma = 0.03$ | | | | $\sigma = 0.3$ | | | |
| orig. poles | Pencil | ESPRIT | MUSIC | orig. poles | Pencil | ESPRIT | MUSIC |
| 0.990 | 0.990 | 0.990 | 1.000 | 0.990 | 0.987 | 0.988 | 1.000 |
| $0.958 \pm 0.248i$ | $0.958 \pm 0.248i$ | $0.958 \pm 0.248i$ | $0.970 \pm 0.239i$ | $0.958 \pm 0.248i$ | $0.965 \pm 0.236i$ | $0.964 \pm 0.239i$ | $0.975 \pm 0.221i$ |
| $0.870 \pm 0.502i$ | $0.870 \pm 0.512i$ | $0.871 \pm 0.512i$ | $0.861 \pm 0.507i$ | $0.870 \pm 0.502i$ | $0.863 \pm 0.511i$ | $0.862 \pm 0.513i$ | $0.880 \pm 0.474i$ |
| $0.662 \pm 0.735i$ | $0.660 \pm 0.737i$ | $0.659 \pm 0.736i$ | $0.712 \pm 0.701i$ | $0.662 \pm 0.735i$ | $0.007 \pm 1.021i$ | $-0.001 \pm 1.012i$ | $-0.034 \pm 0.999i$ |

**Figure 2**

**Heat maps of the wavelet transform**

In Fig. 2, the heat map of the wavelet transform is shown. It follows that yellow region is such that we cannot distinguish two oscillations with close periods. We can recognize 12h and 8h oscillations when the noise is weak. However when the noise is strong (*σ* = 0.3), only the strongest oscillation can be determined. The edge effect is obvious and there are ghost lines e.g. around 15h, that may lead to false estimation.

From these considerations, we conclude that the pencil and ESPRIT methods are robust to noise. This is not the case for MUSIC and CWT.

### Impact of data length

The left-hand side plot of the Fig. 3 shows fit curves using different methods and simulation data (noise standard deviation 0.05) with duration *L* = [30, 50, 100, 200]. The time interval for data collection is 1. Red points indicate simulation data, blue, green and magenta are the fit curves of pencil, ESPRIT and MUSIC algorithms, respectively.

**Figure 3**

**Curves for simulation data**

The right-hand side plot shows poles of oscillations estimated with different methods (noise standard deviation 0.05) with duration *L* = [30, 50, 100, 200]. The time interval for data collection is 1. Black * indicates the original poles of the simulation data, blue, green and magenta are the estimated poles using the pencil, ESPRIT and MUSIC algorithm, respectively. For more accuracy, the poles are also listed in Table 3.

**Table 3.** Poles for different methods

| $L = 30$ | | | | $L = 100$ | | | |
| --- | --- | --- | --- | --- | --- | --- | --- |
| orig. poles | Pencil | ESPRIT | MUSIC | orig. poles | Pencil | ESPRIT | MUSIC |
| 0.995 | 0.896 | -1.043 | 1.000 | 0.995 | 0.994 | 0.994 | 1.000 |
| $0.964 \pm 0.249i$ | $0.778 \pm 0.661i$ | $0.305 \pm 0.000i$ | $0.977 \pm 0.213i$ | $0.964 \pm 0.249i$ | $0.964 \pm 0.249i$ | $0.964 \pm 0.249i$ | $0.969 \pm 0.246i$ |
| $0.863 \pm 0.505i$ | $0.447 \pm 0.000i$ | $0.772 - 0.653i$ | $0.806 \pm 0.591i$ | $0.863 \pm 0.505i$ | $0.863 \pm 0.508i$ | $0.863 \pm 0.508i$ | $0.857 \pm 0.514i$ |
| $0.665 \pm 0.739i$ | $1.093 - 0.329i$ | $1.085 \pm 0.324i$ | $0.456 \pm 0.889i$ | $0.665 \pm 0.739i$ | $0.661 \pm 0.734i$ | $0.659 \pm 0.733i$ | $0.648 \pm 0.761i$ |
| $L = 50$ | | | | $L = 200$ | | | |
| orig. poles | Pencil | ESPRIT | MUSIC | orig. poles | Pencil | ESPRIT | MUSIC |
| 0.995 | 0.995 | 0.995 | 1.000 | 0.995 | 0.995 | 0.995 | 1.000 |
| $0.964 \pm 0.249i$ | $0.964 \pm 0.250i$ | $0.964 \pm 0.250i$ | $0.970 \pm 0.239i$ | $0.964 \pm 0.249i$ | $0.964 \pm 0.249i$ | $0.964 \pm 0.249i$ | $0.972 \pm 0.234i$ |
| $0.863 \pm 0.505i$ | $0.864 \pm 0.511i$ | $0.863 \pm 0.510i$ | $0.824 \pm 0.566i$ | $0.863 \pm 0.505i$ | $0.863 \pm 0.508i$ | $0.863 \pm 0.508i$ | $0.857 \pm 0.514i$ |
| $0.665 \pm 0.739i$ | $0.655 \pm 0.727i$ | $0.652 \pm 0.731i$ | $-0.336 \pm 0.941i$ | $0.665 \pm 0.739i$ | $0.663 \pm 0.737i$ | $0.663 \pm 0.737i$ | $-0.336 \pm 0.941i$ |

### Rate of data collection (sampling frequency)

To investigate the impact of sampling of the underlying continuous-time signal, we generate artificial data with *L* = 50. Then we apply all methods to the original dataset, the half-data set (time collection interval *I* = 2) and third-data set (that is 1, 4, 7, 10… with time collection interval *I* = 3). In Fig. 4, the left-hand side plot below shows heat maps (Y-axis is frequency domain, X-axis is time domain) of simulation data (noise standard deviation 0.05) with duration *L* = [30, 50, 100, 200]. The right-hand side plot shows data fit for the various methods.

**Figure 4**

**Heat maps (left) and fit curves (right)**

#### Conclusion

From the above considerations it follows that decreasing the sampling frequency does not affect the estimation significantly. This means that the data rate collection (sampling frequency) is not an important factor. In contrast, the data length is a crucial factor for all methods.

## Experimental Results: the pencil method applied to gene data

In this section we analyze a small part of the measured data in order to validata some of the aspects of the pencil method and its comparison with the other methods.

**Batch consisting of 171 measurements every 40min**

The results in this case are summarized in Table 4 and Fig. 5.

**Table 4.** Data averaged over all mice

| A | P | T |
| --- | --- | --- |
| 0.1594 | 0.9022 | – |
| 0.0010 | 1.0050 | 1.4483 |
| 0.0017 | 0.9985 | 1.8434 |
| 0.0034 | 0.9956 | 9.8050 |
| 0.0164 | 1.0013 | 23.9361 |
| 0.9239 | 0.9986 | dc |

**Figure 5**

**Plots for averaged data**

**Batch consisting of RER for restrictively fed mice (218 meas. every 40min)** (see Table 5).

**Table 5.** Model parameters for mouse # 1

| Mouse #1 |  |  |
| --- | --- | --- |
| A | P | T |
| 0.0037 | 1.0005 | 4.8275 |
| 0.0116 | 0.9961 | 7.4236 |
| 0.0256 | 0.9993 | 7.9961 |
| 0.0010 | 1.0043 | 20.2774 |
| 0.0817 | 1.0001 | 23.9264 |
| 0.8843 | 1.0001 | dc |

**Figure 6**

**Plots for mouse #1**

Fig. 6 shows the relative approximation error (left pane) and the angle between approximant and error (right pane). Table 6 shows the error and the angles.

**Table 6.** Errors and Angles

| Relative approximation error |  |  |  |  | Angle between approximant & error |  |  |  |  |
| --- | --- | --- | --- | --- | --- | --- | --- | --- | --- |
|  | 3-fit | 5-fit | 7-fit | 9-fit |  | 3-fit | 5-fit | 7-fit | 9-fit |
| Gene 1 | 0.1973 | 0.1276 | 0.1122 | 0.1299 | Gene 1 | 88.72 | 88.65 | 88.66 | 90.46 |
| Gene 2 | 0.2217 | 0.2028 | 0.1669 | 0.1375 | Gene 2 | 88.00 | 89.84 | 87.27 | 86.17 |
| Gene 3 | 0.2801 | 0.3940 | 0.2038 | 0.2112 | Gene 3 | 91.92 | — | 92.25 | 91.54 |
| Gene 4 | 0.2654 | 0.2525 | — | 0.2026 | Gene 4 | 89.82 | 94.18 | — | 92.30 |
| Gene 5 | 0.4296 | 0.3780 | 0.1970 | — | Gene 5 | 84.35 | 86.36 | 89.74 | — |
| Gene 6 | 0.2493 | 0.2563 | 0.1918 | 0.1929 | Gene 6 | 86.94 | 91.78 | 88.39 | 88.78 |
| Gene 7 | 0.1971 | 0.1525 | 0.1475 | 0.1547 | Gene 7 | 89.71 | 88.23 | 88.33 | 90.17 |
| Gene 8 | 0.1914 | 0.1681 | 0.1402 | 0.1619 | Gene 8 | 87.45 | 88.19 | 87.02 | 89.11 |
| Gene 9 | 0.1832 | 0.1913 | 0.1403 | 0.1357 | Gene 9 | 86.36 | 92.63 | 86.64 | 86.68 |
| Gene 10 | 0.2016 | 0.2013 | 0.1874 | 0.2089 | Gene 10 | 86.78 | 87.81 | 86.42 | 89.90 |
| Gene 11 | 0.2637 | 0.2623 | — | 0.2083 | Gene 11 | 92.80 | 91.36 | — | 90.92 |
| Gene 12 | 0.2174 | 0.1681 | 0.2116 | 0.1484 | Gene 12 | 91.20 | 90.18 | 94.12 | 90.59 |
| Gene 13 | 0.3420 | 0.2154 | — | 0.2270 | Gene 13 | 87.25 | 88.50 | — | 91.57 |
| Gene 14 | 0.3140 | 0.2671 | 0.2452 | 0.2034 | Gene 14 | 90.36 | 94.35 | 93.30 | 91.35 |
| Gene 15 | 0.4058 | 0.3374 | 0.3052 | 0.2281 | Gene 15 | 88.15 | 84.41 | 91.66 | 90.31 |

We analyze the relationship among the decomposed oscillations, by calculating the angle among these oscillations for 10 different genes. We set *r* = 9, i.e. the gene signals contain four oscillations **f**_*i*_, *i* = 1,*…*, 4. The approximant is thus 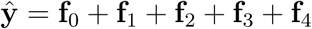. See also Table 7.

**Table 7.** Angle between error vector and approximates

| Gene | $r = 3$ | $r = 5$ | $r = 7$ | $r = 9$ |
| --- | --- | --- | --- | --- |
| Bmal | 89.4040 | 89.0189 | 88.7227 | 89.4645 |
| Clock | 97.5846 | 95.6007 | – | 154.5354 |
| per1 | 87.3120 | 87.0905 | – | 122.6093 |
| per2 | 84.0943 | 84.3410 | 84.2252 | 97.1281 |
| cry1 | 83.6787 | 85.7345 | 83.9466 | – |
| cry2 | 88.0607 | 85.8548 | 85.7156 | 87.9577 |
| rorc | 88.2740 | 87.0592 | 90.5345 | – |
| rora | 92.5359 | – | 90.2449 | 90.3424 |
| rev-erba | 93.4881 | 92.5612 | 91.1162 | 91.4786 |
| reb-rebb | 89.2219 | 89.2972 | 89.0471 | 90.6819 |

From the above tables, we can see that the angle between oscillations is around 90^*°*^ in most situations. So oscillations are nearly orthogonal:

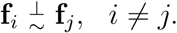

It has actually been shown in [13] that these oscillations are independent of each other.

**Batch consisting of various measurements using mice – 38 min intervals** (see Table 8 and Table 9).

**Table 9.**
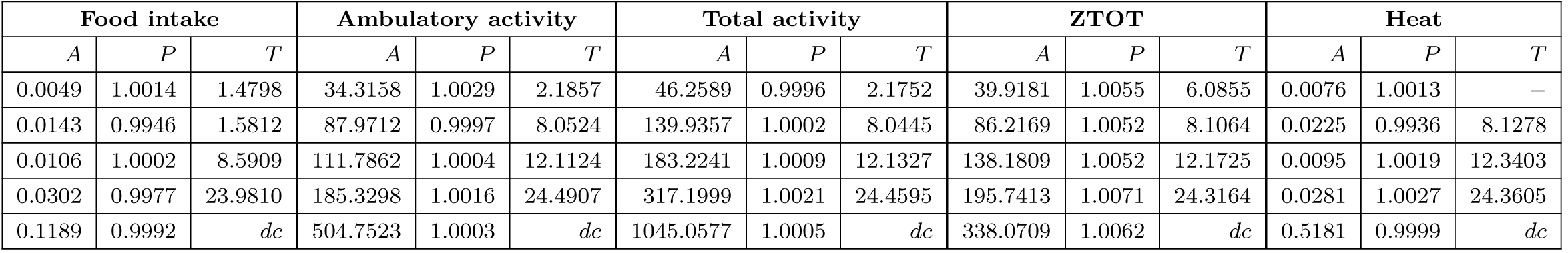
Model parameters for various activities

| Food intake |  |  | Ambulatory activity |  |  | Total activity |  |  | ZTOT |  |  | Heat |  |  |
| --- | --- | --- | --- | --- | --- | --- | --- | --- | --- | --- | --- | --- | --- | --- |
| $A$ | $P$ | $T$ | $A$ | $P$ | $T$ | $A$ | $P$ | $T$ | $A$ | $P$ | $T$ | $A$ | $P$ | $T$ |
| 0.0049 | 1.0014 | 1.4798 | 34.3158 | 1.0029 | 2.1857 | 46.2589 | 0.9996 | 2.1752 | 39.9181 | 1.0055 | 6.0855 | 0.0076 | 1.0013 | — |
| 0.0143 | 0.9946 | 1.5812 | 87.9712 | 0.9997 | 8.0524 | 139.9357 | 1.0002 | 8.0445 | 86.2169 | 1.0052 | 8.1064 | 0.0225 | 0.9936 | 8.1278 |
| 0.0106 | 1.0002 | 8.5909 | 111.7862 | 1.0004 | 12.1124 | 183.2241 | 1.0009 | 12.1327 | 138.1809 | 1.0052 | 12.1725 | 0.0095 | 1.0019 | 12.3403 |
| 0.0302 | 0.9977 | 23.9810 | 185.3298 | 1.0016 | 24.4907 | 317.1999 | 1.0021 | 24.4595 | 195.7413 | 1.0071 | 24.3164 | 0.0281 | 1.0027 | 24.3605 |
| 0.1189 | 0.9992 | $dc$ | 504.7523 | 1.0003 | $dc$ | 1045.0577 | 1.0005 | $dc$ | 338.0709 | 1.0062 | $dc$ | 0.5181 | 0.9999 | $dc$ |

**Table 8.**
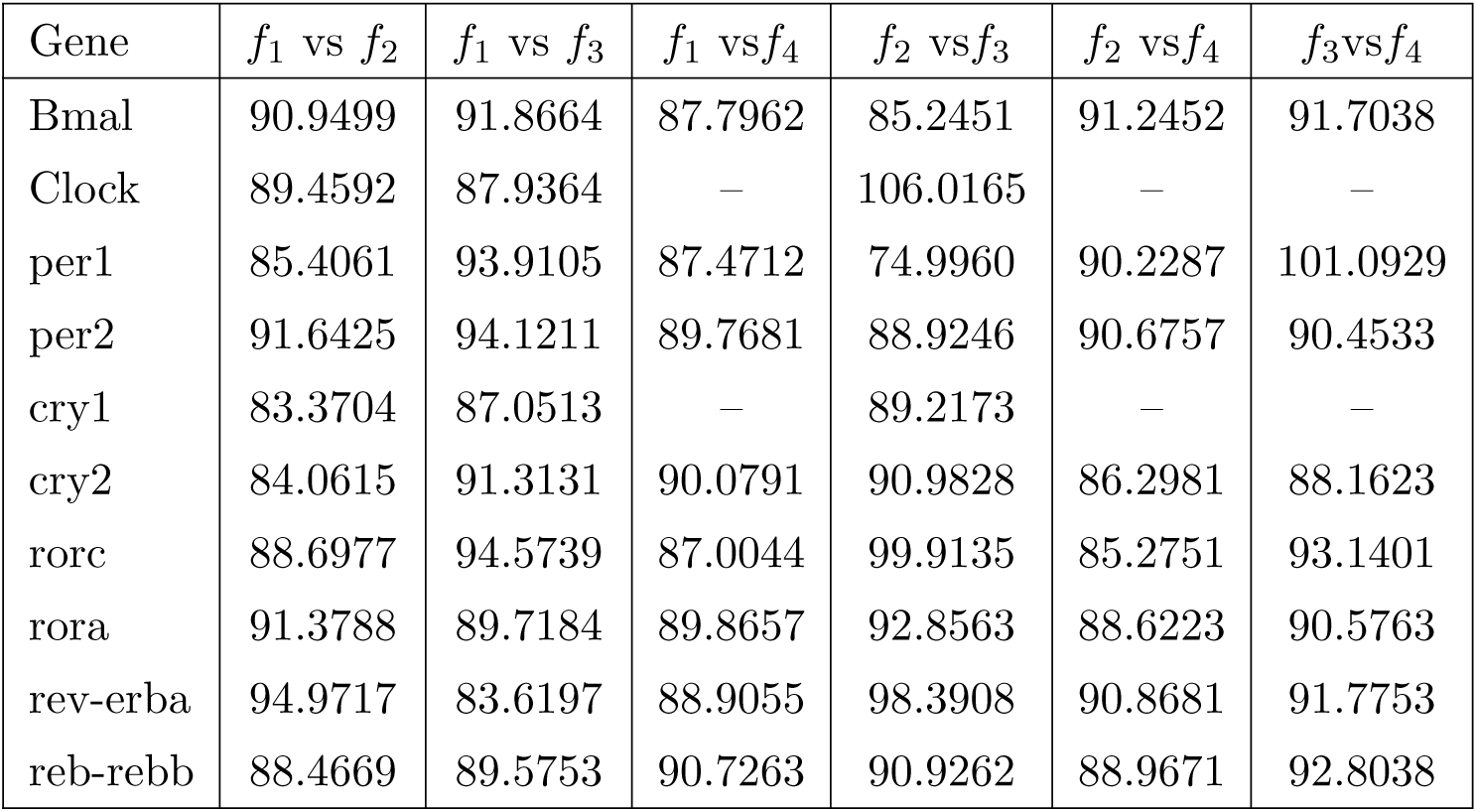
Angle between oscillations

| Gene | $f_1$ vs $f_2$ | $f_1$ vs $f_3$ | $f_1$ vs $f_4$ | $f_2$ vs $f_3$ | $f_2$ vs $f_4$ | $f_3$ vs $f_4$ |
| --- | --- | --- | --- | --- | --- | --- |
| Bmal | 90.9499 | 91.8664 | 87.7962 | 85.2451 | 91.2452 | 91.7038 |
| Clock | 89.4592 | 87.9364 | — | 106.0165 | — | — |
| per1 | 85.4061 | 93.9105 | 87.4712 | 74.9960 | 90.2287 | 101.0929 |
| per2 | 91.6425 | 94.1211 | 89.7681 | 88.9246 | 90.6757 | 90.4533 |
| cry1 | 83.3704 | 87.0513 | — | 89.2173 | — | — |
| cry2 | 84.0615 | 91.3131 | 90.0791 | 90.9828 | 86.2981 | 88.1623 |
| rorc | 88.6977 | 94.5739 | 87.0044 | 99.9135 | 85.2751 | 93.1401 |
| rora | 91.3788 | 89.7184 | 89.8657 | 92.8563 | 88.6223 | 90.5763 |
| rev-erba | 94.9717 | 83.6197 | 88.9055 | 98.3908 | 90.8681 | 91.7753 |
| reb-rebb | 88.4669 | 89.5753 | 90.7263 | 90.9262 | 88.9671 | 92.8038 |

**Figure 7**

**Ambulatory activity: approximation and oscillations**

### Variation of data collection rate

We compare the oscillations using all data (AD), the first half of the data (FHD), the second half of the data (SHD), odd-position data (OD), and even-position data (ED). This is done for a particular set of measurements, but the results are indicative of what happens in general.

Table 10 shows the estimated periods using different part of the data. It follows that the estimation of periods is consistent using AD, FHD, SHD.

**Table 10.** Periods estimated using different parts of the data

|  | AD/h | FHD/h | SHD/h | OD/h | ED/h |
| --- | --- | --- | --- | --- | --- |
| 1 | 24.37 | 23.01 | 24.36 | 24.37 | 24.37 |
| 2 | 12.34 | 12.41 | 12.46 | 11.90 | 12.58 |
| 3 | 8.12 | 8.42 | 7.45 | 8.25 | 8.13 |

## Discussion and comments

### 1. Orthogonality

Recall the definition of angle between signals defined by (6), and let the original vector of measurements for one gene be denoted by **y** *∈* ℝ^*N*^; let also **f**_*i*_,*i* = 0,1,2,3,4, denote the vectors of the DC-component and of the first four fundamental oscillations obtained by means of the pencil reduction method described above. Then the corresponding approximant is ŷ = f_0_ + f_1_ + f_2_ + f_3_ + f_4_.It follows that:

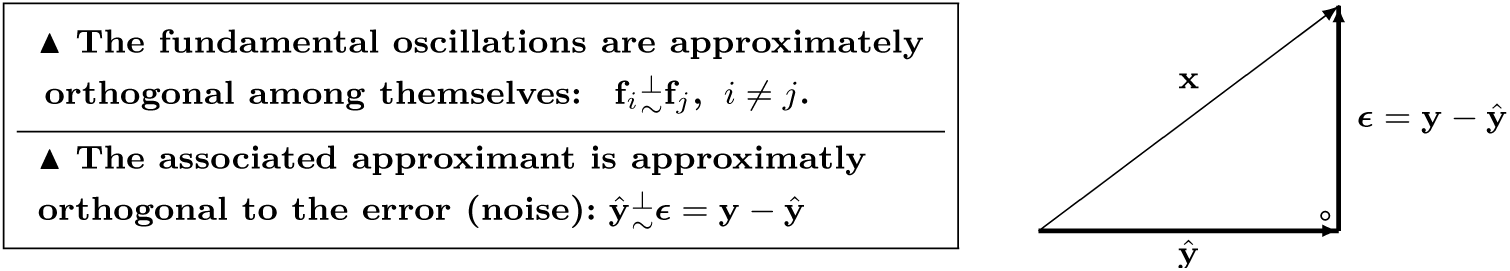

These claims will be justified below.

### 2. Interpretation of orthogonality

Orthogonality means that once an oscillation (e.g. the circadian or the 12h rythm) has been determined, further computations will **not** affect these oscillations. In other words the fundamental oscillations are **independent** of each other.

### 3. Manifestation of orthogonality

As we determine higher-order approximants, i.e. as we add oscillations to the model, the existing ones remain mostly unchanged. Considering the case of the **para probe1** gene, we apply the ESPRIT, LS (Prony’s) and pencil methods. The statistical methods (e.g. ARSER) are not used because being non-parametric they do not allow the choice of the order of fit. ESPRIT and LS are not reliable for large orders of fit, therefore the results for the 24-fit model is not shown. The poles of these three methods are depicted in Tables 11, 12, 13.

**Table 12.** Poles for the LS method

| LS (Prony's method) |  |  |  |
| --- | --- | --- | --- |
| 3 – fit | 5 – fit | 7 – fit | 9 – fit |
| 0.967 | 0.970 | 0.972 | 0.994 |
| 0.363 | $0.435 \pm 0.319i$ | $0.339 \pm 0.354i$ | $0.863 \pm 0.384i$ |
| | $-0.486 \pm 0.366i$ | $-0.517 \pm 0.380i$ | $0.319 \pm 0.863i$ |
| | | 0.363 | $-0.475 \pm 0.745i$ |
| | | | $-0.806 \pm 0.299i$ |

**Table 13.** Poles for the pencil method

| Pencil method |  |  |  |  |
| --- | --- | --- | --- | --- |
| 3-fit | 5-fit | 7-fit | 9-fit | 24-fit (all data) |
| 0.9933 | 0.9932 | 0.9931 | 0.9930 | 0.9915 |
| $0.9436 \pm 0.2734i$ | $0.9449 \pm 0.2730i$ | $0.9446 \pm 0.2742i$ | $0.9447 \pm 0.2747i$ | $0.9489 \pm 0.2843i$ |
| | $0.8609 \pm 0.5132i$ | $0.8659 \pm 0.5086i$ | $0.8672 \pm 0.5068i$ | $0.8729 \pm 0.4812i$ |
| | | $0.3831 \pm 0.9159i$ | $0.3902 \pm 0.9121i$ | $0.3214 \pm 1.1528i$ |
| | | | $-0.9780 \pm 0.3415i$ | $-0.9368 \pm 0.3683i$ |

**Table 11.** Poles for the ESPRIT method

| ESPRIT |  |  |  |
| --- | --- | --- | --- |
| 3 – fit | 5 – fit | 7 – fit | 9 – fit |
| 0.993 | 0.993 | 0.993 | 0.993 |
| $0.939 \pm 0.273i$ | $0.944 \pm 0.272i$ | $0.943 \pm 0.274i$ | $0.944 \pm 0.274i$ |
| | $0.859 \pm 0.509i$ | $0.866 \pm 0.505i$ | $0.866 \pm 0.505i$ |
| | | $0.370 \pm 0.892i$ | $0.374 \pm 0.899i$ |
| | | | $-0.832 \pm 0.213i$ |

### 4. Connection with the Fourier transform

The above method provides an *almost* orthogonal decomposition of a discrete-time signal. The question arises therefore as to whether the same or improved results can be obtained using the Fourier transform and in particular the DFT. Applying the DFT to a length *N* sequence we obtain a decomposition in terms of the *N* **given frequencies or periods**, which are (in decreasing order) 48, 24, 16, 12, 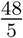, 8, 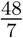, 6, 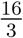, …, 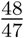. Therefore unless the frequencies of the underlying oscillations are *exactly* among the ones above, the results of the DFT are not useful.

### 5. The least squares (Prony’s) method

This method is not appropriate for cases where the poles are on or close to the unit circle (pure or almost pure oscillations). The figure on the right depicts this fact in the case of the RER data. The conclusion is that while the *matrix pencil method* (red dots) gives oscillatory poles, this is by far not the case with the LS (prony’s) method (green dots).

**Figure 8**

**Comparison between pencil and LS poles**

### 6. Comparison of different methods

(see Table 14).

**Table 14.** Strengths and weaknesses of the various methods

| Method | Parameter Estimation |  |  |  | Estimation Performance |  | Detection of orthogonality |
| --- | --- | --- | --- | --- | --- | --- | --- |
|  | Period | Decay Rate | Amplitude | Phase | Accuracy | Robustness |  |
| DFT | Yes | No | Yes | Yes | Low | Yes | No |
| Wavelet | Yes | Yes | Yes | No | Low | No | No |
| MUSIC | Yes | No | No | No | High | No | No |
| ESPRIT | Yes | Yes | No | No | High | Yes | No |
| Prony (LS) | Yes | Yes | No | No | No | No | No |
| **Pencil** | **Yes** | **Yes** | **Yes** | **Yes** | **High** | **Yes** | **Yes** |

## Final result

We considered a dataset consisting of 18484 genes; transcription is analyzed using the **pencil method** [3], the ESPRIT method, Prony’s method and the three statistical methods. The distribution of the poles follow; recall that the poles of ideal oscillations have magnitude equal to 1.

**Figure 9**

**Results of analysis of 18484 genes using various methods**

Furthermore the DFT and wavelet methods are also not competitive.

The above distributions show that the pencil method has uncovered real oscillations, since the mean of the magnitude of all poles is 1.0058 and the standard deviation is 0.0010. The ESPRIT method follows in terms of discovering oscillations, while the Prony or LS (least squares) method and the three statistical methods give weak results. As explained above the main drawback of the ESPRIT method is that it has nothing to say about the orthogonality of the oscillations, which proves to be a key outcome of the pencil method.

## Concluding remarks and outlook

The matrix pencil method allows the consistent determination of the dominant reduced-order models, thus revealing the fundamental oscillations present in the data. The essence of the matrix pencil method is that it provides a continuous-time tool for treating a discrete-time (sampled-data) problem. The DFT, in contrast, is only a discrete-time tool for treating a discrete-time problem; hence its failure in this setting.

A key consequence of the matrix-pencil approach is the demonstration of orthogonality of the different oscillatory components, in particular the 24-hour and the 12-hour cycles. This points to an independence of these oscillations. This assertion has been subsequently confirmed in the laboratory experiments reported in [13].

This analysis demonstrates the applicability of signal processing methodologies to biological systems and further shows the ability of the matrix pencil decomposition to demonstrate independence of biological rhythms.

